# Insulin at the Intersection of Thermoregulation and Glucose Homeostasis

**DOI:** 10.1101/2023.11.17.566254

**Authors:** Nathan C. Winn, Michael W. Schleh, Jamie N. Garcia, Louise Lantier, Owen P. McGuinness, Joslin A. Blair, Alyssa H. Hasty, David H. Wasserman

**Affiliations:** Department of Molecular Physiology and Biophysics, Vanderbilt University, Nashville, Tennessee, USA; Vanderbilt Mouse Metabolic Phenotyping Center, Nashville, Tennessee, USA; VA Tennessee Valley Healthcare System, Nashville, Tennessee, USA

**Author notes:** **Correspondence:** Nathan C. Winn, Ph.D. Light Hall Rm. 813 Department of Molecular Physiology and Biophysics Vanderbilt University School of Medicine Nashville, TN 37232-0615.

## Abstract

Mammals are protected from changes in environmental temperature by altering energetic processes that modify heat production. Insulin is the dominant stimulus of glucose uptake and metabolism, which are fundamental for thermogenic processes. The purpose of this work was to determine the interaction of ambient temperature induced changes in energy expenditure (EE) on the insulin sensitivity of glucose fluxes. Short-term and adaptive responses to thermoneutral temperature (TN, ∼28°C) and room (laboratory) temperature (RT, ∼22°C) were studied in mice. This range of temperature does not cause detectable changes in circulating catecholamines or shivering and postabsorptive glucose homeostasis is maintained. We tested the hypothesis that a decrease in EE that occurs with TN causes insulin resistance and that this reduction in insulin action and EE is reversed upon short term (<12h) transition to RT. Insulin-stimulated glucose disposal (Rd) and tissue specific glucose uptake were assessed combining isotopic tracers with hyperinsulinemic-euglycemic clamps. EE and insulin-stimulated Rd are both decreased (∼50%) in TN-adapted vs RT-adapted mice. When RT-adapted mice are switched to TN, EE rapidly decreases and Rd is reduced by ∼50%. TN-adapted mice switched to RT exhibit a rapid increase in EE, but whole body insulin-stimulated Rd remains at the low rates of TN-adapted mice. In contrast, whole body glycolytic flux rose with EE. This higher EE occurs without increasing glucose uptake from the blood, but rather by diverting glucose from glucose storage to glycolysis. In addition to adaptations in insulin action, ‘insulin-independent’ glucose uptake in brown fat is exquisitely sensitive to thermoregulation. These results show that insulin action adjusts to non-stressful changes in ambient temperature to contribute to the support of body temperature homeostasis without compromising glucose homeostasis.

**Highlights:**

1. Energy expenditure and insulin-mediated glucose fluxes are reduced in thermoneutral (TN)-adapted mice versus room ‘laboratory’ temperature (RT)-adapted mice.
2. Reduced insulin sensitivity manifests in TN mice regardless of whether they are TN-adapted or short-term transitioned from RT-adapted to TN.
3. TN-adapted mice are resistant to the RT-induced increase in whole-body insulin sensitivity even though metabolic rate is increased.
4. TN-adapted mice switched to RT meets increased thermogenic needs, not by increasing glucose uptake, but by partitioning a greater fraction of glucose from glycogen storage to glycolysis.
5. Brown fat glucose uptake sensitively increases with RT and decreases with TN by an insulin-independent mechanism.

## INTRODUCTION

Energy expenditure (EE) in mammals is sensitively regulated to maintain body temperature homeostasis (1). EE increases to generate heat when ambient temperature falls below thermoneutrality. This increase in metabolic rate is reversible when ambient room temperature is restored to thermoneutral. To meet these changes in EE, substrate oxidation must be dynamic. Insulin is the dominant stimulus of glucose uptake and metabolism in skeletal muscle and adipose tissue. Yet, whether it is involved in the accelerated metabolic rate required to regulate body temperature has not been well defined. It has long since been established that insulin is both critical to maintaining glucose homeostasis and causes a thermogenic response to feeding (2, 3). Low metabolic rate directly relates to insulin resistance – a risk factor for metabolic syndrome and type 2 diabetes (4–6). Conversely, increased EE due to physical activity improves insulin sensitivity and decreases risk factors for metabolic diseases (7). As with physical exercise, intense cold exposure that requires shivering thermogenesis dramatically increases EE for heat generation and associates with increased insulin sensitivity (8–13). The increase in insulin sensitivity with shivering thermogenesis may be comparable to the increase seen with physical exercise.

Body temperature defense during intense cold exposure, like physical exercise, is characterized by sympathetic and adrenomedullary secretion of catecholamines (14, 15). Whether insulin action increases with a mild decrease in environmental temperature that does not require shivering thermogenesis (i.e., a stress response) is unclear. This is particularly relevant in humans, because with modernization, the “burden” of physiological thermoregulation is shared between indoor thermostat-controlled heating and cooling systems that conceptually replicate the body’s temperature control system. Consequently, the physiological challenge created by environmental temperature swings are less extreme and less frequent. Modern societies generally require people spend less time outside for work and recreation (16). The decreased need to accommodate extremes in environmental temperature is associated with a reduction in daily EE (17). This takes on added significance because in addition to temperature regulation, EE is also a major component of body weight regulation, as it is the counterbalance to energy intake. Sustained conditions where EE is less than energy intake leads to weight gain and excess adiposity.

The actions of insulin are a potential intersection for homeostasis of two tightly regulated control systems – arterial glucose and body temperature. The dual role of insulin is significant as challenges to one system may have implications for the second system. For example, ambient temperature may alter how insulin and its counterregulatory hormones maintain glucose homeostasis, create a risk for metabolic disease, or affect insulin dosing for individuals treated with insulin.

The aim of the experiments described herein was to determine the relationship of ambient temperature on the insulin sensitivity of glucose fluxes independent of shivering thermogenesis and the stress of severe decreases in environmental temperature. Short-term and adaptive responses to thermoneutral temperature (TN, ∼28°C) and room (laboratory) temperature (∼22°C) were studied. These temperatures allow for uncomplicated analysis of insulin action because body weights and arterial stress hormone concentrations of chow-fed mice at TN and RT are similar over a ∼4 week adaptive period (18–21). These temperatures also have a practical application to the study of insulin action, glucose metabolism, and EE in mice housed and studied at standard vivaria and laboratory temperatures (∼22°C). Moreover, investigating metabolic phenotypes between RT and TN are more relevant to humans than contrasting RT with more common experimental cold exposures of <15°C (22, 23). Results show that at TN, mice have decreased insulin action and EE regardless of whether mice are TN-adapted or short-term transitioned from RT-adapted to TN. The reciprocal transition from TN-adapted to RT increases EE but unexpectedly without a parallel increase in insulin-stimulated whole body glucose uptake. This higher EE occurs without increasing glucose uptake from the blood, but rather by diverting glucose from glucose storage to glycolysis.

## METHODS

All procedures were approved in advance and carried out in compliance with the Vanderbilt University Institutional Animal Care and Use Committee. Vanderbilt University is accredited by the Association for Assessment and Accreditation of Laboratory Animal Care International.

### Animals and Experimental Design

Male C57BL/6J mice were purchased from Jackson Labs at 11 wks of age and placed on 10% fat diet (Research Diets #D12450B, 3.85 kcal·g^-1^, 70% carbohydrate, 20% protein, 10% fat). At 12 wks of age, mice were transferred to temperature-controlled rooms set at either 22°C or 28°C for 4-6 wks with measurements including body weight, energy homeostasis, body composition, and glucose fluxes (**Fig. 1A**).

**Figure 1.**
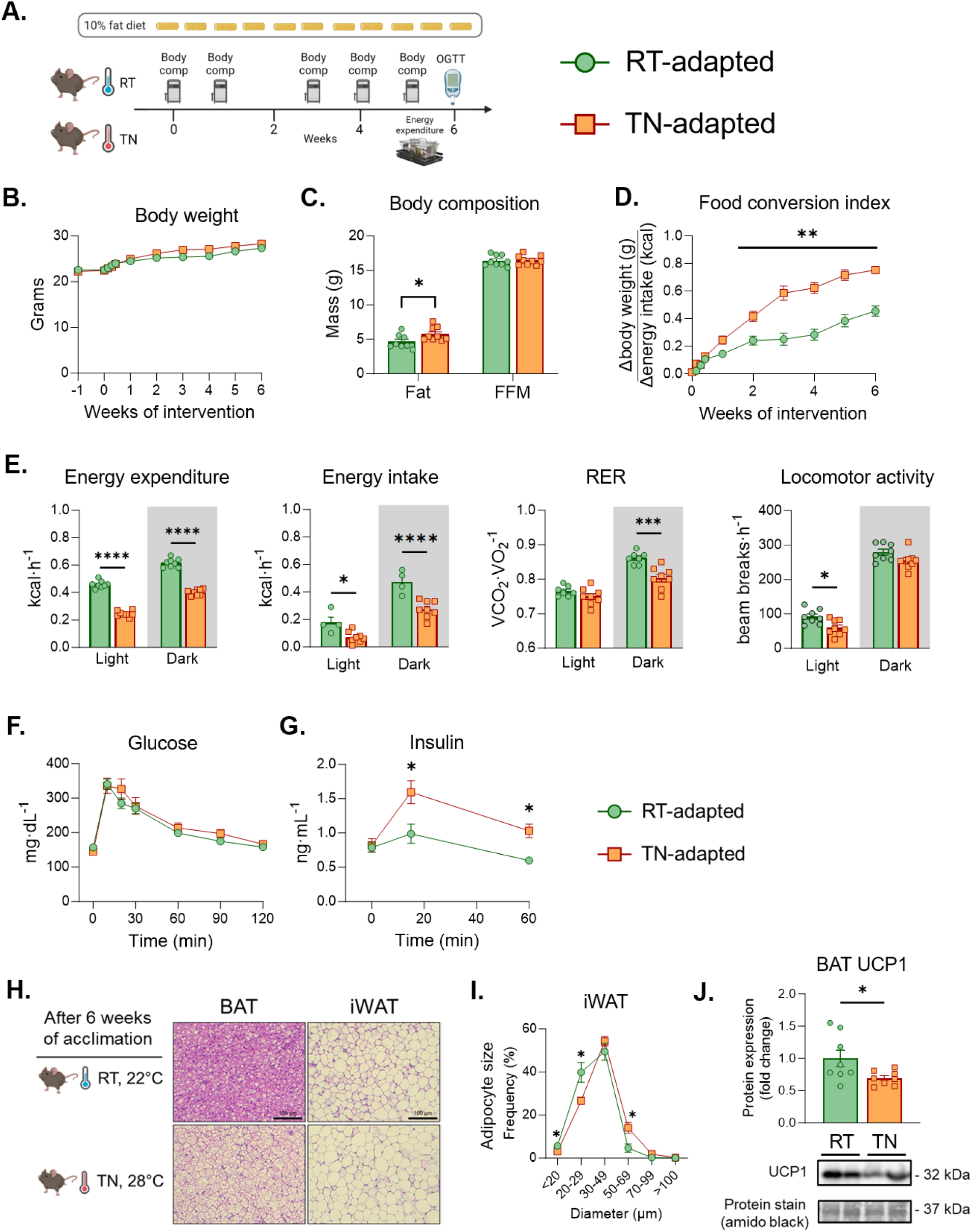
Metabolic phenotype in TN-adapted and RT-adapted mice. **A)** Mice were acclimated to RT or TN for 6 weeks. Glucose tolerance was measured after 5 weeks of acclimation and EE via metabolic cages were performed at week 6. **B)** Body weight was measured over the course of the experiment. **C)** Body composition measurements between RT and TN mice. **D)** Food conversion index was computed as the difference in body weight gain over energy consumed. **E)** EE, energy intake, RER and locomotor activity were collected in real time using metabolic cages with gas analyzers. F) Oral glucose tolerance were performed after 5 weeks of temperature acclimation. **G)** Plasma insulin levels were measured for both groups over the course of 60 minutes after glucose administration. **H**) H&E micrographs of BAT and iWAT. **I**) Adipocyte size distribution in iWAT. **J**) UCP1 immunoblotting in BAT. Panels (B), (D), (F), and (G) two-way ANOVA with repeated measures were run with Tukey adjustment. Students T test was run to compare groups for panel (C). For panel (E), two-way ANOVA with group and photoperiod were run, with Tukey correction. Data are mean ± SE. *p<0.05; **p<0.01; ***p<0.001; ****p<0.0001

### Body composition

Mouse body fat mass, fat-free mass (FFM), and free water were measured by a nuclear magnetic resonance whole-body composition analyzer (Bruker Minispec).

### Glucose tolerance testing

Oral glucose tolerance tests (OGTT) were performed at the temperatures at which the mice were housed (i.e., 22°C or 28°C). After a 5-hour fast, basal blood glucose was measured (-10 min) in samples obtained from a tail cut followed by oral glucose gavage of 3.5 g/kg fat-free mass. Blood glucose was sampled at 15, 20, 30, 60, 90, and 120 min after administration using a hand-held glucometer (Bayer Contour Next EZ meter). Approximately 20 μl of blood was obtained at t=0, 15, and 60 min for measurement of insulin.

### Vehicle infusion and hyperinsulinemic-euglycemic (insulin) clamps

Catheters were surgically placed in a carotid artery and jugular vein for sampling and infusions, respectively, one week before clamp procedures (24). Mice were transferred to a 1.5 L plastic container without access to food 5 hour prior to the start of an experiment (**Fig. S1A**). Glucose clamps were conducted as described previously (24). Mice were neither restrained nor handled during clamp experiments. [3-^3^H]glucose was primed and continuously infused from t=-90min to t=0 min (0.06 µCi·min^-1^). The insulin clamp was initiated at t=0 min with a continuous insulin infusion (4 mU·kg^-1^·min^-1^) and variable glucose infusion initiated and maintained until t=155 min. The glucose infusate contained [3-^3^H]glucose (0.06 µCi·µl^-1^) to minimize changes in plasma [3-^3^H]glucose specific activity. Arterial glucose was monitored every 10 min to provide feedback to adjust the glucose infusion rate (GIR) as needed to maintain euglycemia. Erythrocytes were infused at a rate calculated to compensate for blood withdrawal over the duration of the experiment. [3-^3^H]glucose specific activity was determined at -15 min and -5 min for the basal period, and every 10 min between 80 to 120 min for the clamp period to assess glucose disappearance (Rd) and endogenous glucose production (EndoRa). Whole-body glycolytic rate was determined by the ^3^H_2_O formation rate, and glucose storage was calculated as the difference between Rd and glycolysis, as detailed previously (25). A 13 µCi intravenous bolus of 2-[^14^C]-deoxyglucose ([^14^C]2DG) was administered at 120 min to determine the glucose metabolic index (Rg), an index of tissue-specific glucose uptake. Blood samples were collected at 122, 125, 130, 135 and 145 min to measure plasma [^14^C]2DG. At 145 min, mice were euthanized and tissues immediately harvested and freeze-clamped. For vehicle infusion studies, a continuous saline infusion (infusion rate = 1µl·min^-1^) was started at t=0 and procedures were identical to those described for the insulin clamp. Detailed methodology is available via VMMPC webpage https://vmmpc.org.

### Chemical Analyses

Plasma non-esterified fatty acids (NEFA) (Wako HR series NEFA-HR, FUJIFILM Wako Diagnostics U.S.A) were determined via colorimetric assay according to the manufacturer’s instructions. Endogenous plasma insulin was determined by radioimmunoassay (Millipore Cat. # PI-13K), as described (26). The assay uses ^125^I-labled insulin and a double antibody technique to determine plasma insulin. The cross-reactivity between human and mouse insulin is 100% in the radioimmunoassay used. Exogenous insulin was determined with use of a human-specific insulin antibody (no. 80-INSHU-E01.1; Alpco). All fasting blood samples were collected after a 5-hour fast. Glycogen assay was performed according to Chan and Exton (27).

### Indirect calorimetry and ambulatory activity

Respiratory gases, locomotor activity, and feeding were determined using the Promethion Metabolic Analyzer (Sable Systems, North Las Vegas, NV). Rates of EE were calculated from 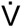O_2_ and 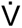CO_2_ using the Weir equation [*EE* (*kcal · h*^−1^) = 60 · (0.003941 ·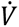*O*_2_) + (0.001106 · 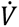*CO*_2_)]. After 4 weeks of acclimation to either RT or TN, mice had metabolic sensors and activity sensors fit atop their home cages. Mice were maintained at the temperature at which they were acclimated. A 24-hour run-in period was used to allow mice to adjust to the presence of sensors in the home cage. Baseline ‘acclimated’ measurements were recorded for 72 hours (0-72 h). After 72 hours, the temperature in the chamber was either increased to TN or decreased to RT in a subset of mice and measurements were collected from 72-144 h. Non-protein substrate oxidation from carbohydrate [*Carbohydrate oxidation* (*mg* · *min*^−1^) = (4.344 · 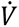*CO*_2_) − (3.061 · 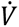*CO*_2_)] and fat [*Fat oxidation* (*mg* · *min*^−1^) = (1.695 · 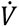*O*_2_) − (1.707 · 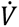*O*_2_)] were estimated according to Jeukendrup and Wallis (28). Carbohydrate and fat oxidation rates were converted to kcal·h^-1^ using the energy equivalent 4.0 kcal·g^-1^ for carbohydrate and 9.75 kcal·g^-1^ for fat (28). A fixed rate of protein oxidation was assumed based on the duration of the indirect calorimetry measurements (29).

### Untargeted metabolomics

Metabolomics analyses were performed by the Vanderbilt Center for Innovative Technology. After a 5-hour fast, plasma, liver, heart, gastrocnemius muscle, inguinal adipose tissue (iWAT), epididymal adipose tissue (eWAT), and brown adipose tissue (BAT) were quickly excised, and flash frozen in liquid nitrogen. Tissue processing was performed as previously described (30). Briefly, samples were collected using a Thermo Scientific Q Exactive HF (LC Hybrid Quadrupole Orbitrap MS/MS) instrument. Data were aligned using Progenesis QI 2.0 software and analyzed using the MetaboanalystR package (R4.3.1) (31). Between group differences were compared by paired samples t-test from the global metabolome detected across all tissues (206 metabolites) and were adjusted to a <5% false discovery rate (Benjamini-Hochberg method).

### Statistical Analyses

Student’s t-tests were run for between group comparisons. If data did not follow a Gaussian distribution, non-parametric Mann-Whitney tests were used to determine statistical significance. In experiments that contained more than two groups, one-way analysis of variance (ANOVA) or two-way ANOVA models were conducted with pairwise comparisons using Tukey or Sidak correction. Brown-Forsythe correction was applied to groups with unequal variance. The EE ANCOVA analysis was conducted using the NIDDK MMPC (www.mmpc.org) Energy Expenditure Analysis website (http://www.mmpc.org/shared/regression.aspx). Data are presented as mean ± standard error (SE). An adjusted p value of <0.05 was used to determine significance.

## RESULTS

### Metabolic phenotype in TN-adapted and RT-adapted mice

Mice acclimatized to either RT (22°C) or TN (28°C) for 6 weeks (**Fig. 1A**) have similar body weight gain (**Fig. 1B**), despite a small increase in fat mass in TN mice (∼1.5 g, **Fig. 1C**). There are no differences in FFM. An index of food conversion (the ratio of body weight gain to kcal consumed) is 33% higher in TN-adapted compared to RT-adapted mice (**Fig. 1D**). EE and energy intake are decreased in TN mice during both light and dark photoperiods (**Fig. 1E**). The respiratory exchange ratio (RER) is also lower during the dark photoperiod in TN mice (**Fig. 1E**). Locomotor activity is not different between RT-adapted and TN-adapted mice during the dark photoperiod, whereas TN-adapted mice show less physical activity during the light photoperiod. The latter may be explained by increased foraging during the light photoperiod and subsequent food consumption by the RT-adapted mice. An OGTT was performed to determine whether long-term reduction in EE at TN worsens glucose tolerance. Fasting glucose and fasting insulin are not different between groups (**Fig. 1F&G**). Following oral glucose, the blood glucose peak and subsequent decline are not different between RT-adapted and TN-adapted mice (**Fig. 1F**). However, TN-adapted mice required a greater glucose-stimulated arterial insulin to maintain normal glucose tolerance compared to RT-adapted mice (**Fig. 1F&G**). This is consistent with lower insulin sensitivity in TN-adapted mice. BAT and iWAT are densely innervated by sympathetic nerves and are sensitive to changes in sympathetic tone (8, 32–34). As expected, BAT and iWAT from TN-adapted mice display larger lipid droplets and adipocytes, respectively than RT-adapted mice (**Fig. 1H&I**). Uncoupling protein 1 (UCP1), an important protein in non-shivering thermogenesis (35), is also lower in BAT from TN-adapted mice (**Fig. 1J**). No notable morphological differences between RT-adapted and TN-adapted mice are observed for eWAT or liver (**Fig. S2A**).

### Metabolome signatures in TN-adapted versus RT-adapted mice

To further characterize how ambient temperature modifies metabolism, metabolomic analyses were performed in arterial plasma and several insulin-sensitive tissues from postabsorptive mice. Plasma metabolomics reveal 51 differentially expressed metabolites (**Fig. 2A**). Pathway analysis shows an enrichment in glycerophospholipid metabolism and non-polar aromatic amino acid biosynthesis (**Fig. 2B**). TN-adapted have higher circulating phospholipids (**Fig. 2C**); whereas one of the most decreased metabolites in TN-adapted mice is glycerol-3-phosphate – a cellular regulator of cytosolic NAD+ (36) and a synthetic precursor to fibroblast growth factor-23 (37). The BAT metabolome is highly sensitive to environmental temperature, with 75 differentially expressed metabolites (**Fig. 2F & Fig S3**). Amino acid biosynthesis and ketone body synthesis/degradation are among the upregulated pathways (**Fig. 2E**). L-methionine is increased in TN-adapted mice, which is inversely associated with UCP1 content and BAT lipid content (38). In contrast to BAT and plasma, other tissues show fewer differences in metabolites between RT-adapted and TN-adapted (**Fig. 2G** & **Fig. S2**). No single metabolite was up or down across all tissues in RT-adapted or TN-adapted. However, ophthalmic acid, which increases when cellular glutathione turnover is high is a common metabolite decreased in TN-adapted skeletal muscle, heart, iWAT, and BAT (39). Metabolic intermediates of taurine metabolism are also increased in all TN-adapted tissues except iWAT. Taurine metabolism may increase as a defensive response to inflammation and oxidative stress (40, 41).

**Figure. 2.**
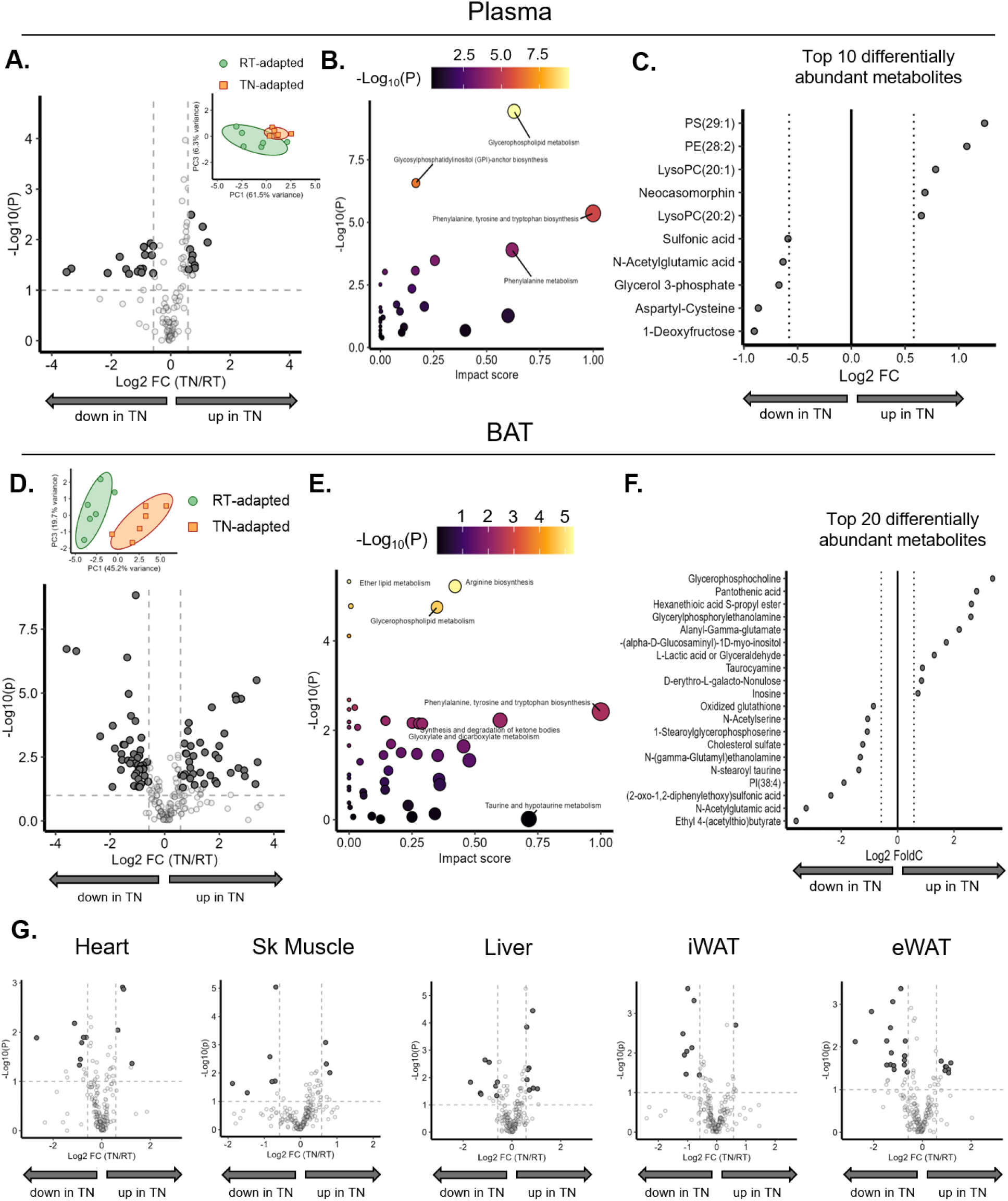
Metabolome signatures in TN-adapted versus RT-adapted mice. After 6 weeks of temperature acclimation, mice were euthanized and plasma and tissues extracted and flash frozen after a 5 hour fast. Untargeted metabolomics were performed in the plasma and BAT compartments. **A**) Volcano plot of differentially abundant plasma metabolites, with PCA plot as the inset. **B**) KEGG pathway score of differentially abundant metabolites. **C**) Top 10 differentially abundant plasma metabolites significantly increased in RT and TN, respectively. **D**) Volcano plot of differentially abundant BAT metabolites, with PCA plot presented as an inset. **E**) KEGG pathway score of differentially abundant metabolites. **F**) Top 20 differentially abundant BAT metabolites significantly increased in RT and TN, respectively. **G**) Volcano plots for heart, skeletal muscle, liver, iWAT, and eWAT. Student t tests were used to determine between group differences for adipocyte size bins Data are mean ± SE; n=6/group. *p<0.05.

### Glucose fluxes during short-term transition of RT-adapted mice to TN (RT→TN)

RT-adapted mice underwent euglycemic vehicle or insulin clamps, while respiratory gases were determined in a parallel cohort of animals. RT→TN mice were compared to RT-adapted mice in which temperature remained constant (RT→RT) (**Fig. 3A**). The acute increase in temperature leads to an immediate reduction in EE across photoperiods (**Fig. 3B**). Body weight remains unchanged (**Fig. 3C**). Carbohydrate oxidation is decreased in RT→TN during the dark photoperiod, while fat oxidation is lower during both light and dark periods (**Fig. 3D&E**).

**Figure. 3.**
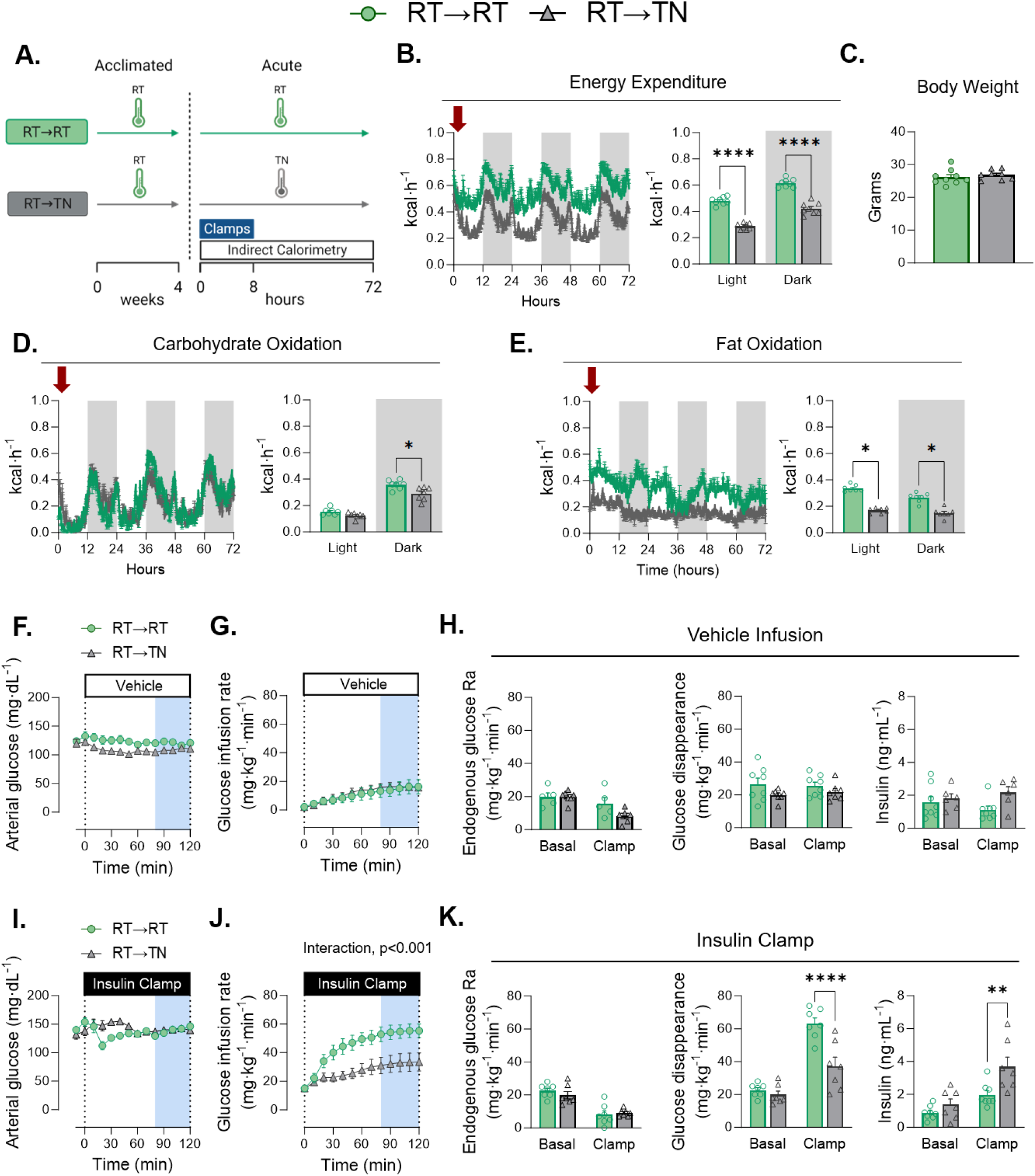
Glucose fluxes during short-term transition of RT-adapted mice to TN (RT→TN). **A)** The experimental model is shown in which mice were housed at RT for 4 weeks. At the beginning of the fasting period prior to clamping, the mice either remained in their temperature environment or transitioned to the opposing temperature. **B)** The environmental temperatures were monitored over the course of the experiment. **C)** EE of mice acclimated to RT or transitioned to novel TN. **D**) Body mass was not different between groups. [E-G, vehicle infusion]. **E**) Arterial glucose was administered and monitored during the GIR. **F)** Glucose infusion rate was monitored for each group. **G)** The rate of endogenous glucose appearance was measured for each group. Glucose flux was measured between groups over 100 minutes. Insulin levels were measured in all groups during clamping. [H-J, insulin clamps]. **H)** Glucose was administered and monitored during insulin infusion. **I**) GIR during clamped insulin infusion. **J**) The rate of endogenous glucose appearance was measured for each group. Glucose flux was measured between groups over 100 minutes. Insulin levels were measured in all groups during clamping. **K**) Glucose disappearance **L**) NEFA levels plotted against insulin concentrations. Student t tests were run to determine group differences for panels C and D. Panels E,F,H,I two-way ANOVA with repeated measures were run with Tukey adjustment. Panels G and J, two-way ANOVA with group and condition as factors were run with Tukey adjustment. Panel K and L, simple linear regression was used to test differences in slopes between groups. Data are mean ± SE. *p<0.05, **p<0.01, ****p<0.0001

We next tested whether the dynamics of EE and substrate oxidation are accompanied by corresponding changes in basal and insulin-mediated glucose fluxes. During the vehicle infusion, a low GIR is necessary to maintain arterial glucose constant as the fast duration progresses during the experiment (**Fig. 3F&G**). GIRs are not different between RT→TN and RT→RT mice (**Fig. 3G**). Similarly, there are no differences between EndoRa, Rd, or plasma insulin during the basal or steady state vehicle infusion (**Fig. 3H**). These data show that with vehicle infusion, whole-body glucose fluxes are not affected by a dynamic increase in ambient temperature.

By design, arterial glucose is clamped at the same glucose concentrations in RT→TN and RT→RT mice (**Fig. 3I**). RT→TN reduces insulin-stimulated GIR by nearly 50% (**Fig. 3J**). The attenuation in GIR is due to a decrease in Rd, with no differences in EndoRa (**Fig. 3K**). This decrease in Rd occurs despite a significant increase in arterial insulin with RT→TN during the insulin clamp (**Fig. 3K**). Human insulin was infused during the insulin clamp creating similar concentrations of exogenous insulin in each cohort (**Table 1**). Plasma NEFAs are not different between RT→TN and RT→RT mice at basal and fell similarly during insulin-stimulated conditions (**Table 1**). No differences in plasma norepinephrine or epinephrine were found between groups (**Table 1**). Together, these data show that acute reduction in EE evoked by an increase in environmental temperature corresponds to a decrease in insulin-stimulated but not basal glucose fluxes.

**Table 1.** Blood chemistry.

| **Table 1 – Blood chemistry** |  |  |  |  |  |  |
| --- | --- | --- | --- | --- | --- | --- |
|  | RT→RT | RT→TN | p value | TN→TN | TN→RT | p value |
| Norepinephrine (pg·mL<sup>-1</sup>) | 457.4 ± 49 | 516.6 ± 82 | p=0.53 | 397.8 ± 69 | 410.2 ± 69 | p=0.90 |
| Epinephrine (pg·mL<sup>-1</sup>) | 168.6 ± 25 | 135.5 ± 12 | p=0.25 | 171.1 ± 24 | 153.4 ± 13 | p=0.53 |
| Basal NEFA (mmol·L<sup>-1</sup>) | 1.03 ± 0.14 | 0.76 ± .14 | p=0.20 | 1.30 ± 0.22 | 1.01 ± 0.13 | p=0.27 |
| Clamp NEFA (mmol·L<sup>-1</sup>) | 0.24 ± 0.03 | 0.19 ± 0.03 | p=0.19 | 0.29 ± 0.02 | 0.38 ± 0.04 | p=0.08 |
| Exogenous insulin levels (ng·mL<sup>-1</sup>) | 2.6 ± 0.09 | 2.5 ± 0.07 | p=0.99 | 2.6 ± 0.06 | 2.9 ± 0.21 | p=0.10 |
| Hematocrit (%) | 39.1 ± 1.7 | 40.0 ± 0.9 | p=0.65 | 41.3 ± 1.2 | 39.4 ± 1.1 | p=0.27 |

A bolus of [^14^C]2DG was injected intravenously to determine tissue-specific Rg during vehicle and insulin clamps (**Fig. 4**). Compared with RT→RT, RT→TN reduces BAT Rg during vehicle and insulin clamps by ∼75 and ∼65%, respectively (**Fig. 4A & B**). No differences in Rg between RT→RT and RT→TN are detectable for skeletal muscle or white AT during vehicle infusion or insulin clamps. However, during the insulin clamp cardiac muscle Rg is decreased in RT→TN compared to RT→RT mice (**Fig. 4B**). Insulin-mediated Rg is computed as the difference between insulin clamp Rg and vehicle infusion Rg. Insulin-mediated cardiac Rg is lower in RT→TN mice compared to RT→RT, whereas insulin-mediated BAT Rg is not significantly decreased (**Fig. 4C**). These data suggest that transition to TN reduces BAT Rg by an insulin-independent mechanism, whereas the reduction in cardiac Rg is insulin-dependent.

**Figure 4.**
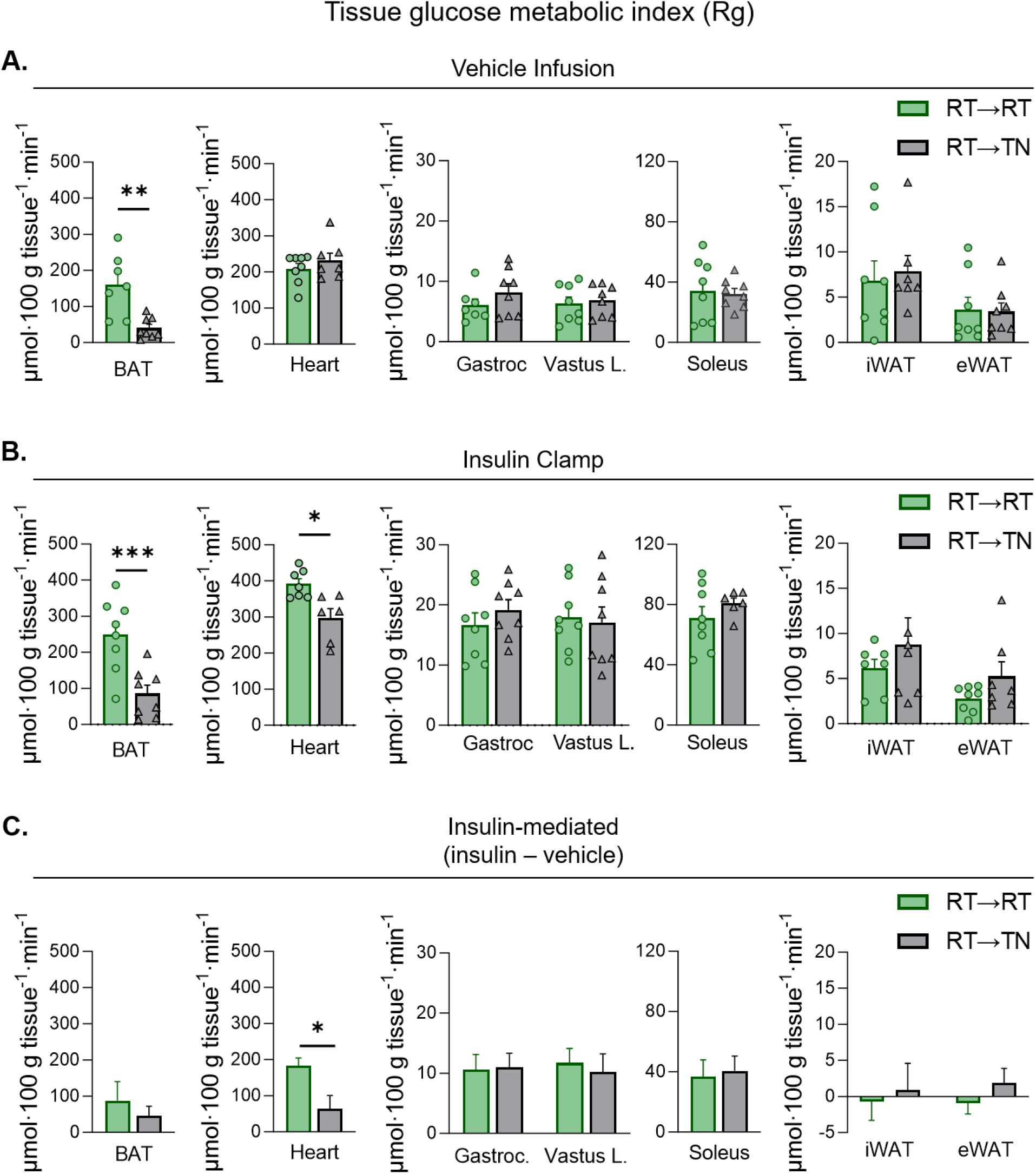
Non-insulin and insulin-mediated tissue glucose metabolic index during short-term transition from RT-adapted to TN (RT→TN). [^14^C]2-deoxyglucose was infused as a bolus at t=120 of each respective clamp. Blood was collected frequently for 25 minutes to determine the rate of disappearance using exponential decay. Tissues were rapidly excised and snap frozen for isotopic enrichment. Tissue Rg between RT→RT and RT →TN during **A**) vehicle and **B**) insulin clamps. **C**) Insulin-stimulated tissue Rg was computed as the mean differences in tissue Rg between insulin and vehicle infusions. The variance between vehicle and insulin clamps were calculated using the standard error of the difference. Statistical significance was determined using a t-distribution table with critical t-values and corresponding degrees of freedom. Data are mean ± SE; n=6-9/group. p<0.05 was used to reject the null hypothesis. *p<0.05, **p<0.01, ***p<0.001

### Glucose fluxes during short-term transition of TN-adapted mice to RT (TN→RT)

Next, we tested the hypothesis that increased EE caused by transition from TN-adapted to RT required an increase in insulin-mediated glucose fluxes. TN→TN and TN→RT groups were compared (**Fig. 5A**). TN→RT leads to a marked increase in EE compared to TN→TN (**Fig. 5B**). Mice in each condition had similar body weights (**Fig. 5C**). Carbohydrate oxidation is markedly increased during the light and dark photoperiods of TN→RT mice. Fat oxidation is higher in TN→RT during the light photoperiod, but not the dark cycle (**Fig. 5E**). These data show that the short-term transition from TN to RT results in an immediate increase in EE that is primarily due to oxidation of carbohydrates.

**Figure 5.**
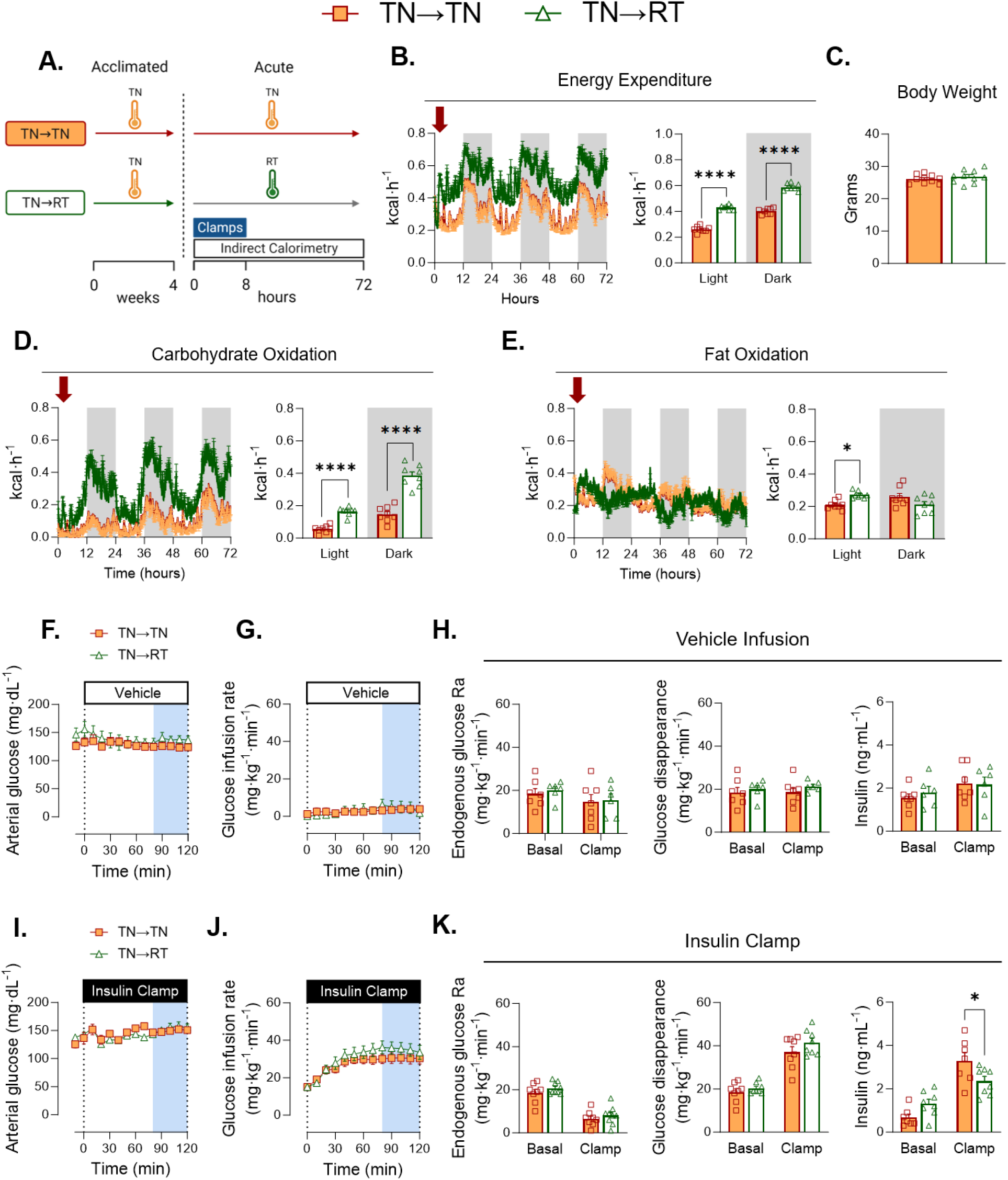
Glucose fluxes during short-term transition of TN-adapted mice to RT (TN→RT). **A**) The experimental model is shown in which mice were housed at either TN for 4 weeks. At the beginning of the fasting period prior to clamping, the mice either remained in their temperature environment or transitioned to the opposing temperature. **B)** The environmental temperatures were monitored over the course of the experiment. **C) EE** of mice acclimated to RT or transitioned to novel TN. **D**) Body mass was not different between groups. [E-G, vehicle infusion]. **E**) Arterial glucose was administered and monitored during the GIR. **F)** Glucose infusion rate was monitored for each group. **G)** The rate of endogenous glucose appearance was measured for each group. Glucose flux was measured between groups over 100 minutes. Insulin levels were measured in all groups during clamping. [H-J, insulin clamps]. **H)** Glucose was administered and monitored during insulin infusion. **I**) GIR during clamped insulin infusion. **J**) The rate of endogenous glucose appearance was measured for each group. Glucose flux was measured between groups over 100 minutes. Insulin levels were measured in all groups during clamping. **K**) Glucose disappearance **L**) NEFA levels plotted against insulin concentrations. Student t tests were run to determine group differences for panels C and D. Panels E, F, H, and I two-way ANOVA with repeated measures were run with Tukey adjustment. Panels G and J, two-way ANOVA with group and condition as factors were run with Tukey adjustment. Panel K and L, simple linear regression was used to test differences in slopes between groups. Data are mean ± SE; n=7-9/group. *p<0.05, ****p<0.0001

Steady state arterial glucose and GIR are not different between TN→TN and TN→RT mice during the vehicle infusion (**Fig. 5 F&G**). Similarly, no differences in EndoRa, Rd, or insulin concentrations are detected during the euglycemic vehicle infusion (**Fig. 5H**). Steady state arterial glucose is achieved during the insulin clamp in all groups (**Fig. 5I**). TN→RT does not increase GIR above TN→TN during an insulin clamp (**Fig. 5J**). Moreover, EndoRa and Rd are also not augmented TN→RT, despite a ∼50% increase in EE. Steady state clamp insulin concentrations are 39% lower in TN→RT versus TN→TN mice (**Fig. 5K**). Exogenous human insulin concentrations are not different between cohorts (**Table 1**). Similarly, no differences in basal or clamp NEFAs are found and circulating catecholamines are comparable between conditions (**Table 1**). Taken together, TN→RT have a robust increase in EE that is accompanied by a large increase in carbohydrate oxidation, but do not exhibit the increase in whole-body insulin-mediated glucose fluxes typically seen at RT.

While TN→RT did not improve whole-body Rd, BAT Rg is increased ∼5-fold and ∼3-fold during vehicle infusions and insulin clamps, respectively (**Fig. 6A & B**). In addition, during the insulin clamp, iWAT Rg is ∼2-fold higher in TN→RT than TN→TN mice (**Fig. 6B**). Interestingly, there is a decrease in vastus lateralis and soleus Rg in TN→RT mice compared to TN→TN during insulin but not during vehicle infusions, with gastrocnemius showing a similar but non-significant trend **(Fig. 6B**). Insulin-mediated vastus lateralis and soleus Rg is lower in TN-RT compared to TN→TN while iWAT Rg is elevated (**Fig. 6C**). Insulin-mediated BAT Rg is not different between TN→RT and TN→TN (**Fig. 6C**).

**Figure 6.**
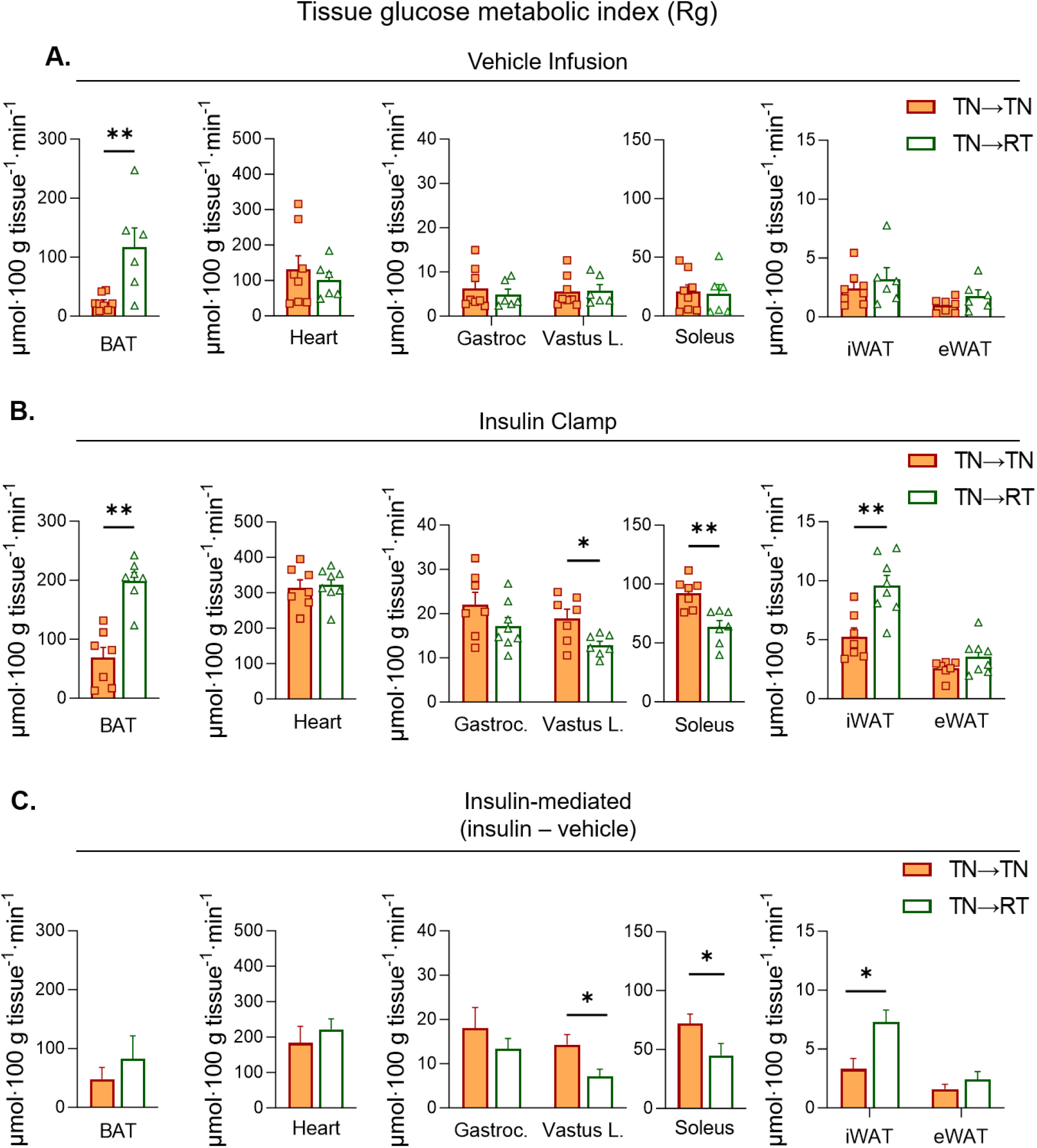
Non-insulin and insulin-mediated tissue glucose metabolic index during short-term transition from TN-adapted to RT (TN→RT). [^14^C]2-deoxyglucose was infused as a bolus at t=120 of each respective clamp. Blood was collected frequently for 25 minutes to determine the rate of disappearance using exponential decay. Tissues were rapidly excised and snap frozen for isotopic enrichment. Tissue Rg between TN→TN and TN →RT during **A**) vehicle and **B**) insulin clamps. **C**) Insulin-stimulated tissue Rg was computed as the mean differences in tissue Rg between insulin and vehicle infusion. The variance between vehicle and insulin clamps were calculated using the standard error of the difference. Statistical significance was determined using a t-distribution table with critical t-values and corresponding degrees of freedom. Data are mean ± SE; n=6-9/group. p<0.05 was used to reject the null hypothesis. *p<0.05, **p<0.01

### Whole-body glucose partitioning during short-term temperature transitions

A reduction in insulin-stimulated glucose storage is a hallmark of insulin resistance (4–6). We quantified the glycolytic rate from the rate of accumulation of ^3^H_2_O released from [3-^3^H]glucose at the triose phosphate step of glycolysis (**Fig. 7A&D**). Glucose storage is the difference between Rd and glycolysis. During vehicle infusions, rates of glycolysis and glucose storage are not different between groups (**Fig. 7B&E**). Insulin increases the rate of glycolysis and glucose storage in RT→RT, whereas only glycolytic flux is significantly enhanced by insulin in RT→TN. (**Fig. 7B**). Skeletal muscle glycogen content is lower in RT→TN mice, while liver glycogen is not different (**Fig. 7C**). In TN→TN mice, insulin increases glucose storage and glycolysis (**Fig. 7E**). The insulin-mediated increase in glucose storage is lost in TN→RT mice (**Fig. 7E**, black bars) as a twofold greater fraction of glucose is diverted to glycolysis compared to TN→TN mice (**Fig. 7E**, blue bars). No differences in skeletal muscle or liver glycogen were found between TN→RT and TN→TN mice (**Fig. 7F**). Glycolytic flux during insulin-stimulation is directly associated with EE (**Fig. 7G**), whereas glucose storage did not correlate with EE (**Fig. 7H**). Overall, the adaptation to acute RT meets increased thermogenic needs, not by increasing Rd, but by partitioning a greater fraction of glucose from glucose storage to glycolysis.

**Figure 7.**
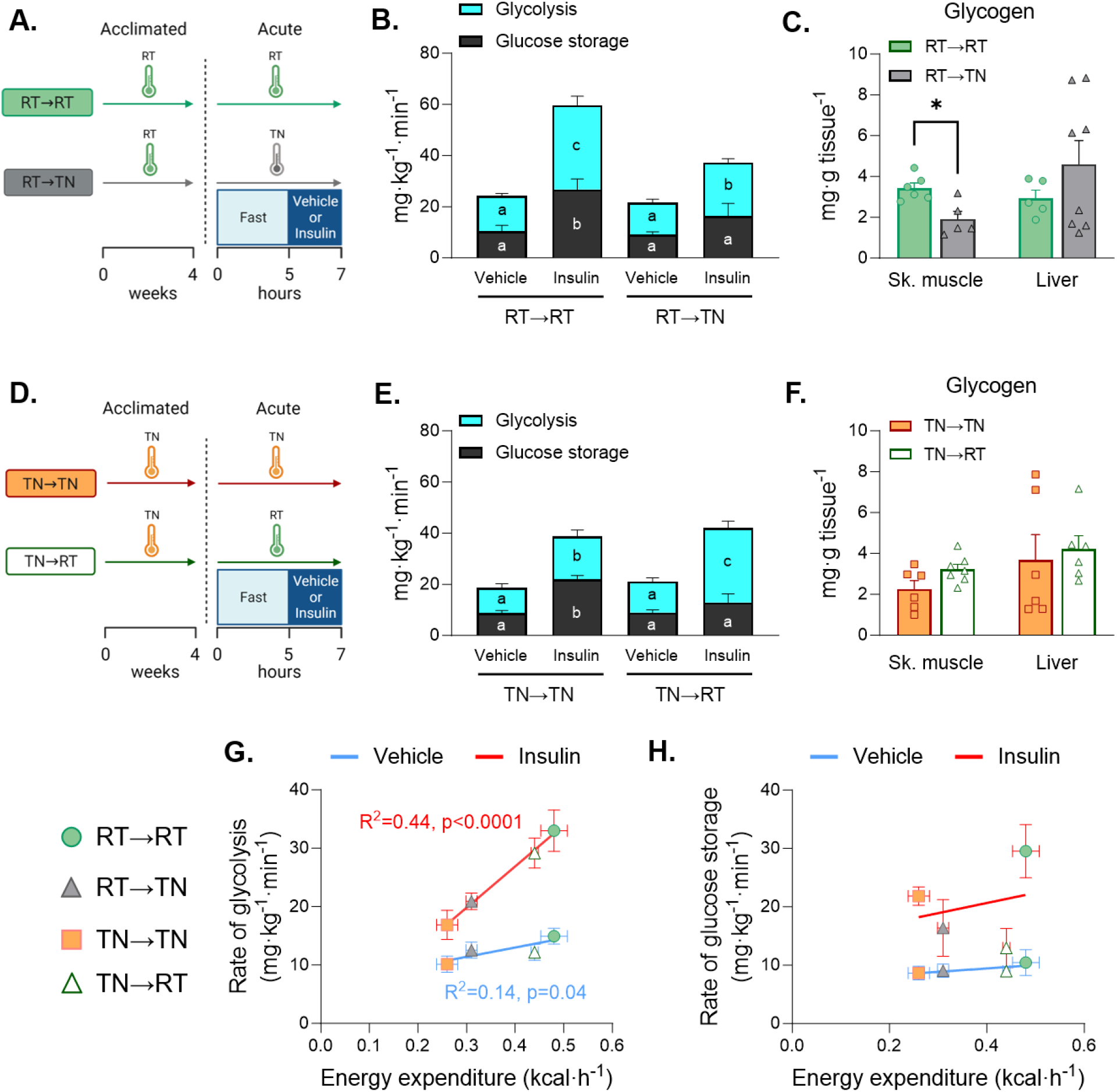
Whole-body glucose partitioning during short-term temperature transitions. Mice acclimated to either RT or TN were short-term exposed to a novel temperature **A**) TN and **B**) RT or remained at their acclimated temperature during a euglycemic vehicle or insulin clamp. The rate of glycolysis and glucose storage were quantified and compared between **C**) RT→TN and RT→RT or **D**) TN→RT and TN→TN conditions. Glycogen was measured and compared between **E**) RT→TN and RT→RT or **F**) TN→RT and TN→TN. **G**) The relationship between the rate of glycolysis and EE was plotted for all temperature conditions. **H**) The rate of glucose storage and EE was plotted for all temperature conditions. For regression plots, blue lines represent vehicle conditions and red lines reflect insulin stimulated conditions. For panels C and D, two-way ANOVA with treatment and temperatures as factors. The ANOVAs were run for glycolysis and glucose storage separately. Histograms with distinct letters are statistically significantly different from one another. Black letters correspond to differences within glycolysis comparisons and white letters denote differences within glucose storage. Independent samples T tests were run for panels E and F for skeletal muscle and liver, respectively. Simple linear regression was used to determine differences in slopes for panels G and H. Data are mean ± SE; n=6-9/group.

## DISCUSSION

Mechanisms that maintain core body temperature are highly conserved in endotherms (1). In response to deviations in environmental temperature, core body temperature remains constant through a balance of heat production and heat dissipation. The mechanism of heat production requires an increase in metabolism, broadly known as thermogenesis. Metabolism is at the intersection between the regulation of glucose homeostasis and temperature homeostasis. Insulin action on metabolism is essential for glucose homeostasis and has been implicated in feeding-induced thermogenesis (2, 3). The mechanism by which insulin is able to participate in the regulation of two highly conserved homeostatic processes without compromising either remains to be elucidated. In this study, we tested the hypothesis that a decrease in metabolic rate that occurs in TN-adapted mice causes insulin resistance and that this reduction in insulin action and EE is reversed upon short-term transition to RT. RT was selected as the low temperature because it does not result in overt stress, seen at lower temperatures, as arterial catecholamine concentrations are not increased (Table 1). As hypothesized, whole-body insulin action is decreased at TN regardless of whether mice are TN-adapted or transitioned short-term from RT to TN. Remarkably, TN-adapted mice do not increase whole-body insulin action during the transition to RT, even though EE transitions rapidly. Moreover, in contrast to RT-adapted mice, which rapidly modulate fat oxidation in response to changes in EE demand, TN-adapted mice rely on increases in carbohydrate oxidation to support the dynamic increase in EE in the transition to RT. This is paralleled by a decrease in insulin-stimulated whole-body glucose storage, which is a hallmark feature of insulin resistant states (4–6, 42). Glucose storage as glycogen requires that glucose is transported across the plasma membrane and then phosphorylated by a hexokinase to form glucose-6-phosphate. Glucose-6-phosphate allosterically stimulates glycogen synthase and glucose storage (43). We show that skeletal muscle glucose Rg is decreased during transition from TN-adapted to RT. Thus, less flux of blood borne glucose to glucose-6-phosphate occurs. Since skeletal muscle, due to its mass, is the primary site of insulin-stimulated glucose storage, a decrease in skeletal muscle Rg may contribute to reduced whole-body glucose storage. Whole-body glycolytic flux is quantitatively increased during TN→RT. Glycolysis is not compromised but continues unabated to adjust to changing temperature. Results suggest that glucose storage only occurs once the energy demand of thermoregulation has been met by glycolysis and metabolism of other substrates. In RT-adapted mice, the metabolic demand is met by both carbohydrate and fat substrates; whereas, TN-adapted mice have a bias towards carbohydrate oxidation.

Basal whole-body glucose fluxes are not different between RT and TN despite pronounced differences in EE. This holds true regardless of whether mice are adapted or dynamically switched to these temperatures. This suggests that a non-stressful environmental temperature shift does not disrupt fasting glycemic control. This also implies that long-term adaptations are not required to maintain tight control of postabsorptive glycemia in the presence of marked changes in EE. Fasting BAT glucose uptake toggles with environmental temperature so that it is elevated at RT and reduced at TN, regardless of whether mice are temperature adapted. Thermogenic adipose tissue, including BAT and beige AT, contributes to heat production via the actions of UCP1, intracellular creatine cycling, and possibly glutamine cycling (8, 44, 45). TN-adapted mice have lower UCP1 and increased lipid expansion in BAT consistent with low thermogenic activity. Furthermore, the relative abundances of metabolic intermediates involved in heat-generating processes are lower in TN-adapted mice (Fig. 1). This includes oxidized glutathione and ophthalmic acid which are increased during the elevated oxidative state of BAT thermogenesis (46, 47). In contrast, intermediates related to inflammatory pathways and collagen remodeling are increased in TN mice, including glycylproline which is a substrate for prolidase – an enzyme linked to collagen turnover (Fig. 2 and (48)). Interestingly, partial loss of prolidase activity in BAT causes local fibrosis and inflammation (48). The increase in these pathways in TN BAT is hypothesized to support cellular remodeling required for increased lipid accumulation (49). Thermogenic fat is insulin-sensitive and has been proposed to be a major site of substrate disposal including glucose (50). Despite its high sensitivity to insulin the increase in BAT glucose uptake in the presence of RT is insulin-independent, regardless of whether they are RT-adapted or not.

To our knowledge, this is the first study to directly characterize the differences between standard laboratory temperature and TN conditions on insulin-dependent and insulin-independent glucose fluxes in the mouse. This is significant considering that mice are typically housed and studied at temperatures that best accommodate investigators (RT) but below mouse thermoneutrality. There are reports that living below thermoneutral temperature may even increase longevity (51). This may be related to improved insulin action at temperatures below thermoneutrality. In this regard, evidence shows that a 25% reduction in arterial insulin increases longevity in mice (52). Experimentally, it may be advantageous to study mice at TN when increases in EE or insulin action are anticipated. EE and insulin action are reduced at TN, making increases easier to resolve. Furthermore, our results show that the metabolic phenotype is most evident when mice are challenged with a provocative stimulus such as insulin stimulation. It may not be feasible to house animals at TN. These studies show that it may be sufficient to perform experiments at TN, without prior TN housing, to obtain conditions that may be advantageous for specific purposes. In addition to amplifying some regulatory mechanisms, long-term housing at TN can be useful as it accelerates the development of cardiovascular and metabolic disease in the mouse (53–55).

A strength of the experimental design is that it allowed for the measurement of basal and insulin-mediated tissue-specific glucose fluxes in the ‘unstressed’ mouse. This was accomplished by measuring glucose fluxes during vehicle infusions and insulin clamps. Questions emerge from the novel findings of these experiments. Insulin-stimulated whole-body Rd remains low in TN-adapted mice ∼8 h after transition to RT even though EE has increased. Despite the lower Rd, glycolysis becomes accelerated and glucose storage is selectively reduced. The molecular mechanisms that mediate the partitioning of glucose in TN-adapted mice remain to be defined. It also remains to be determined how long TN adaptations persist after transition to RT. It would be informative to know the on and off kinetics for both adaptation and reversal of temperature-induced changes in metabolism. This study examined male mice and did not address the possibility of sexual dimorphisms. There are reasons to speculate that females respond differently since they are more insulin sensitive than males after accounting for differences in FFM (56). Increased sympathetic outflow in response to a decrease in temperature stimulates adipose tissue lipolysis to make substrate available for heat production (57, 58). However, the temperature differential tested in this work did not cause differences in arterial NEFAs at baseline or steady state insulin clamp conditions. In addition, circulating catecholamine concentrations were not different between temperature conditions. The possibility of differences at the adipocyte sympathetic synaptic clefts cannot be ruled out. Previous studies show mixed results for the role of differences in sympathetic nerve activity between mice at RT and TN (59, 60).

In aggregate, this study defines the insulin-dependent and insulin-independent regulation of metabolism in response to modest deviations in environmental temperature. Mechanisms for maintaining body temperature homeostasis are such that glucose homeostasis is preserved. It is not lost on the authors that the results have implications for the majority of studies which are conducted at room temperature. Housing and laboratory temperatures are variables to consider in the design of metabolic research *in vivo*. We identify a mismatch between metabolic rate and insulin-stimulated glucose uptake, which is due to impaired glucose storage. Glucose is preferentially channeled to glycolysis to support EE. These studies show that insulin action is at the intersection of thermoregulation and glucose homeostasis. Insulin action adjusts to non-stressful changes in ambient temperature to contribute to the support of body temperature homeostasis without compromising glucose homeostasis.

## ACKNOWLEDGMENTS

We acknowledge the following Vanderbilt University (VU) and Vanderbilt University Medical Center (VUMC) core facilities: VUMC Hormone Assay & Analytical Services Core (NIH DK059637 and DK020593), VU Metabolic Mouse Phenotyping Center [VMMPC (NIH DK059637; www.vmmpc.org)], NIDDK MMPC Energy Expenditure Analysis webpage [(http://www.mmpc.org/shared/regression.aspx) DK076169 and DK115255], Center for Innovative Technology (CIT), and Translational Pathology Shared Resource (NCI/NIH Cancer Center Support Grant 5P30 CA68485-19). This work was supported in part by grants to DHW (DK054902, DK050277), AHH (IK6BX005649) and NCW (DK136926). We thank Alec Rodriguez for mouse husbandry and technical assistance. We thank Tasneem Ansari, Staci Bordash, Teri Doss, Alicia Kellarakos, Carlo Malabanan for their assistance with surgical procedures and clamp studies. We also thank Dr. E. Matthew Morris for intellectual input.

## AUTHOR CONTRIBUTIONS

NCW conceived and designed research, performed experiments, interpreted results, and drafted the manuscript. MWS, performed experiments, interpreted results, and reviewed/edited the manuscript. LL, designed research, interpreted results, and reviewed/edited the manuscript. OPM, interpreted results, designed research, and reviewed/edited the manuscript. JNG, performed experiments and reviewed/edited the manuscript. JAB, performed experiments and reviewed/edited the manuscript. AHH, interpreted results and reviewed/edited the manuscript. DHW, conceived and designed research, interpreted results, and reviewed/edited the manuscript. NCW is the guarantor of this work and, as such, had full access to all the data in the study and takes responsibility for the integrity of the data and the accuracy of the data analysis.

---

**Figure S1.**
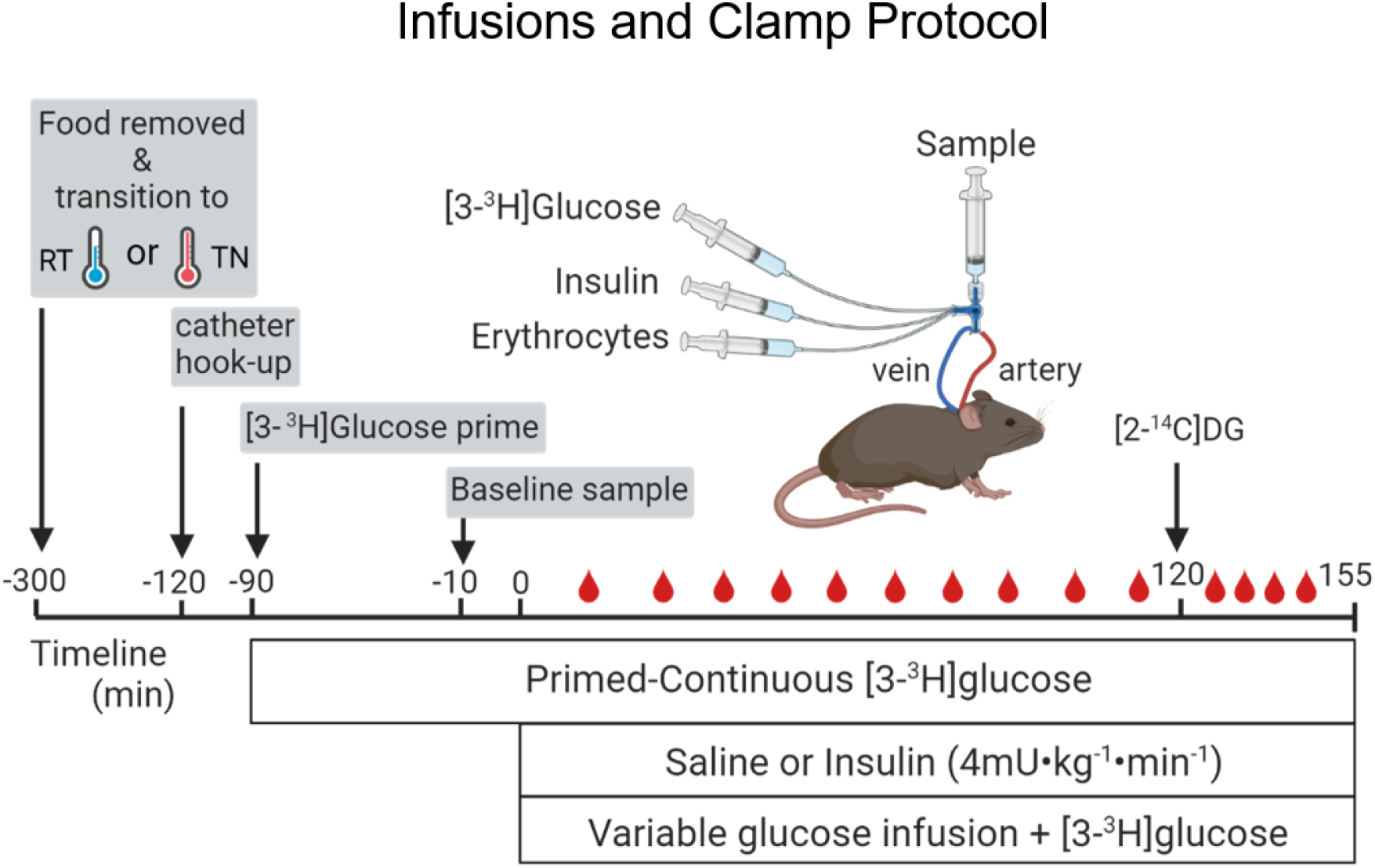
Experimental vehicle and insulin clamp schematic description.

**Figure S2:**
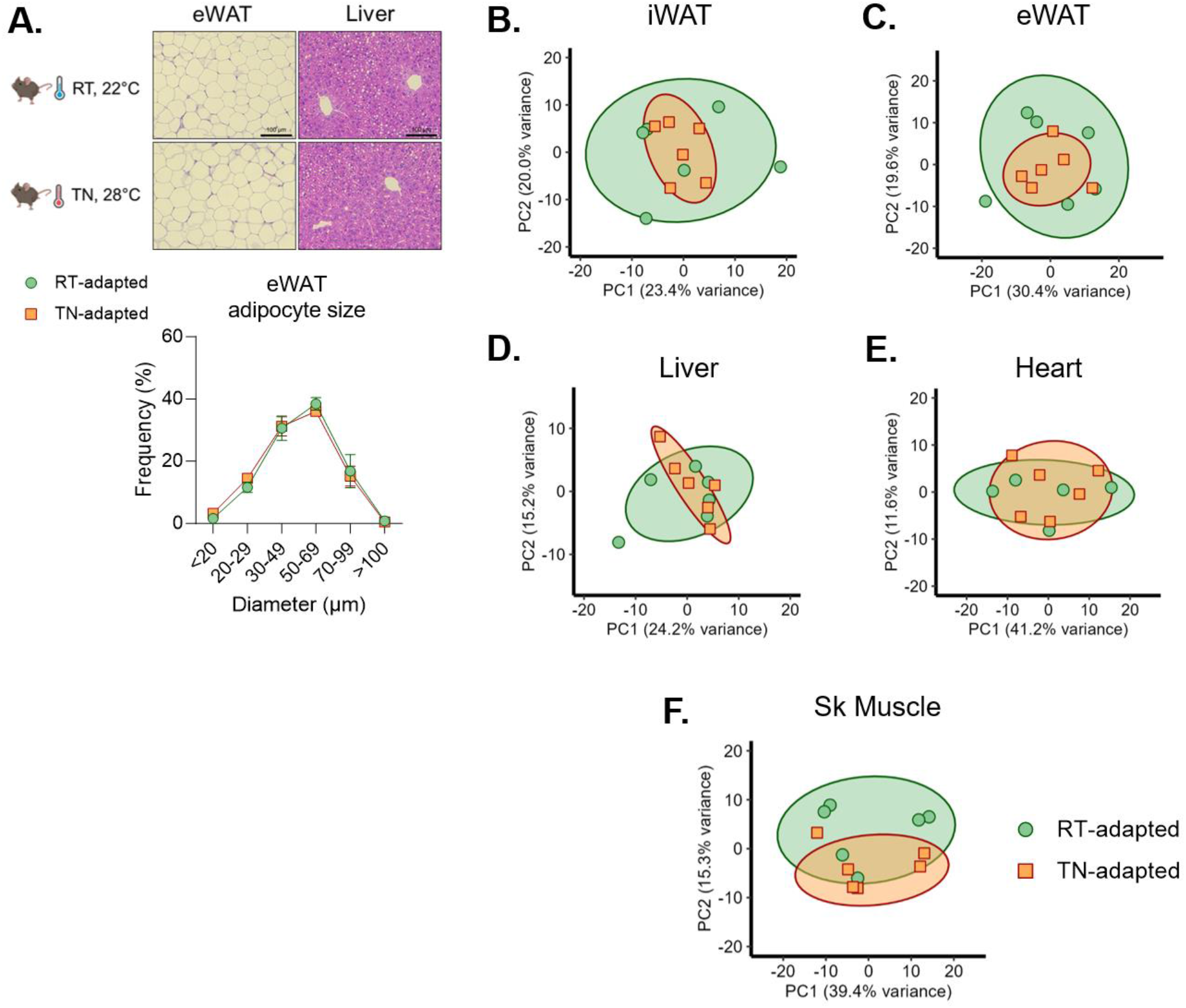
Mice were adapted to RT or TN for 6 weeks. **A)** eWAT and liver micropgraphs were generated. Adipocyte size was calculated via ImageJ and presented as frequency per bin size. After a 5 h fast, plasma, liver, heart, gastrocnemius muscle, inguinal adipose tissue (iWAT), epididymal adipose tissue (eWAT), and brown adipose tissue (BAT) were quickly excised and flash frozen in liquid nitrogen from mice acclimated to RT or TN. Tissues were processed and analyzed using HILIC negative ion mode mass spectrometry (LC MS). Spectral intensities were median normalized and Iog10 transformed. PCA plots were generated via R software package in **B)** iWAT, **C)** eWAT, **D)** liver, **E)** heart, and **F)** skeletal muscle. n=6/group.

**Figure S3:**
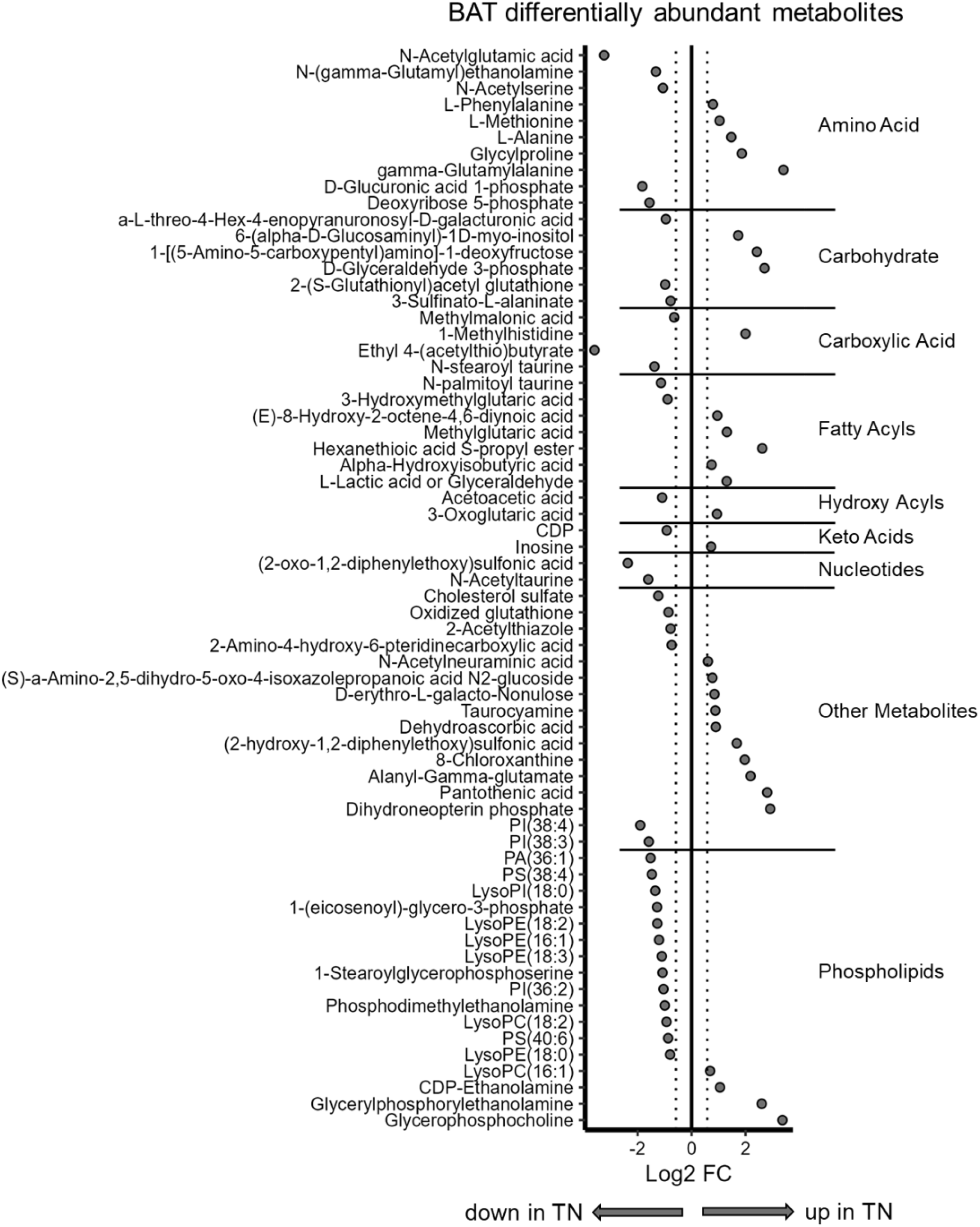
Mice were adapted to RT or TN for 6 weeks. Differentially expressed BAT metabolites are presented relative to RT-adapted control. Metabolites are clustered by class and presented as Log2 fold difference. Significance was accepted if FDR<O.05.

